# Macromolecular crowding limits growth under pressure

**DOI:** 10.1101/2021.06.04.446859

**Authors:** Baptiste Alric, Cécile Formosa-Dague, Etienne Dague, Liam J. Holt, Morgan Delarue

## Abstract

Cells that grow in confined spaces eventually build up mechanical compressive stress. This growth-induced pressure (GIP) decreases cell growth. GIP is important in a multitude of contexts from cancer[1–3], to microbial infections[4], to biofouling, yet our understanding of its origin and molecular consequences remains limited. Here, we combine microfluidic confinement of the yeast Saccha-romyces cerevisiae[5],with rheological measurements using genetically encoded multimeric nanoparticles (GEMs)[6] to reveal that growth-induced pressure is accompanied with an increase in a key cellular physical property: macromolecular crowding. We develop a fully calibrated model that predicts how increased macromolecular crowding hinders protein expression and thus diminishes cell growth. This model is sufficient to explain the coupling of growth rate to pressure without the need for specific molecular sensors or signaling cascades. As molecular crowding is similar across all domains of life, this could be a deeply conserved mechanism of biomechanical feedback that allows environmental sensing originating from the fundamental physical properties of cells.

---

Cells in every kingdom of life can proliferate in spatially-limited environments. In metazoans, tissues have physical boundaries[7]. In plants, roots sprout into a solid ground[8, 9]. In microbes, substrate adhesion physically limits colony expansion[10–12]. To proliferate in confinement, cells must push on the boundaries of their environment and neighboring cells, leading to development of compressive forces that translate, at the multicellular scale, into the buildup of a mechanical growth-induced pressure, hereafter denoted GIP. GIP decreases cell growth and division of all organisms: bacteria, fungi, plants or mammals[1–3, 5, 13, 14]. However, the mechanisms that control proliferation under GIP remain unknown. In particular, it is unclear whether growth reduction is due to specific signaling pathways, or is a necessary consequence to changes of the physical properties of the cells.

Some signaling pathways have been associated with survival or division under GIP[15, 16], but it remains unclear if these pathways affect growth *per se*. For example, mutants in the SCWISh network, composed of the Cell Wall Integrity pathway and signaling from *Ste11* through *Msb2/Sho1* proteins tend to lyse due to mechanical instabilities associated with budding, but their ability to develop GIP is unperturbed[16].

On the other hand, mechanical perturbations to cells also influence fundamental physical parameters. One such parameter is macromolecular crowding, which relates to the high packing fraction of macromolecules in the cell, and can decrease biochemical reaction rates due to decreased effective diffusion[17–20]. However, the role of crowding in response to mechanical stress in general, and GIP in particular, has been largely overlooked.

In this Letter, we investigated the relationship between growth-induced pressure, macromolecular crowding and cell growth in the budding yeast *S. cerevisiae*. Our results are best explained by a model in which the rates of intracellular osmolyte production and macromolecular biogenesis are intrinsically coupled. To develop GIP, osmolytes and macromolecules are produced, while cell expansion is limited, causing the cell interior to become crowded, leading to a biophysical feedback that limits cell growth.

We used microfluidic elastic chambers as a model confining 3D environment (Fig. 1a, more details in Fig. S1)[21]. After filling the chamber, cells pushed against their neighbors and onto their surroundings. Cells were continually fed through microchannels to prevent nutrient depletion and enable switching of media. After 10 hours of confined growth, the elastic chamber was quite deformed, almost doubling in volume. This deformation was used to measure the amount of growth-induced pressure, GIP, developed by the cells[5, 16]. We posited that, under confinement, GIP resulted from an increase in intracellular osmotic pressure, which was balanced not only by the cell wall but also by the surrounding effective elasticity of the other cells and the PDMS chamber.

**Figure 1.**
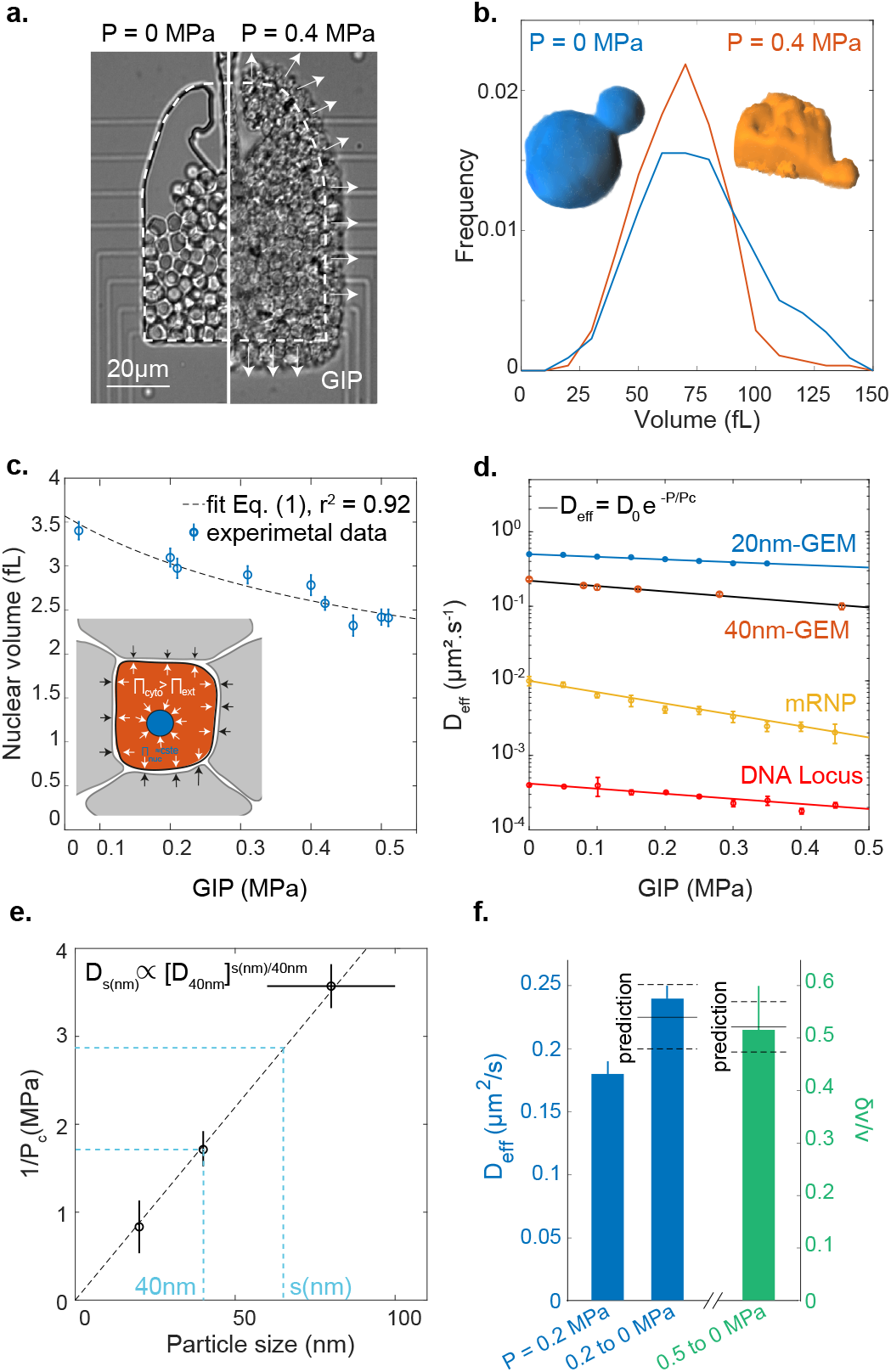
Confined growth leads to the intracellular accumulation of osmolytes and macromolecules. **a.** Confined growth leads to the build-up of growth-induced pressure (GIP), measured by the deformation of the PDMS chamber. **b.** Cell volume distribution under GIP. Insets: representative 3D reconstructions of a non-compressed cell and a cell at 0.4 MPa. Both cells have a similar volume ~ 65 fL. **c.** Nuclear volume decreases under GIP. Dashed line: fit of nuclear volume as a function of GIP assuming constant nuclear osmotic pressure Eq. (1) (r^2^ = 0.92). **d.** Diffusivities of various particles and a DNA locus all decrease exponentially as a function of GIP. Solid black curve is the model prediction for 40nm-GEMs r^2^ = 0.98. **e.** The characteristic pressure (*P_c_*) of the exponential dependence is inversely proportional to cytosolic particle size. **f.** After sudden pressure relaxation, effective diffusion rises quickly (< 1 minute) to control (uncompressed) values and cell volume increases (*δv*) due to stored osmotic pressure. Predicted values are indicated. The diffusion data fall within 7% of the prediction, while the volume data fall within 2%. In all data, values are mean ±standard error of the mean, N ≥ 3 independent biological replicates.

Remarkably, cell size did not decrease as GIP increased (Fig. 1b), even though cells became highly deformed (Inset Fig. 1b). The deformation of cells was a consequence of compressive forces. These forces can originate from an increase in the intracellular osmotic pressure that, due to confinement, applies forces to the chamber and to surrounding cells, thereby deforming them, like inflating balloons inside a box (see Inset Fig. 1c). Strikingly, we observed a strong reduction in nuclear volume (Fig. 1c) and, as a result, the nuclear-to-cell volume ratio was perturbed. This is distinct from osmotic stress which leads to a proportional reduction of the nuclear volume, keeping the nuclear/cytoplasmic volume ratio constant as in [22] (Fig. S2).

We can subdivide osmolytes into two classes, small and large, that we operationally define by their ability to freely diffuse across the nuclear pore, a cutoff value of ~ 3 nm hydrodynamic radius[23]. The concentration of small osmolytes is dominated by ions and metabolites such as glycerol, while large osmolytes are macromolecules such as proteins, ribosomes and mRNA. The decrease in nuclear volume under osmotic stress is indicative of an increase in the concentration of cytoplasmic macromolecules. The changes in nuclear volume under GIP suggested that the concentration of cytoplasmic macromolecules was also increasing under GIP. In agreement, our data were best fit assuming that these two osmolytes (small osmolytes and macromolecules) were increasing proportionally. Assuming that nuclear osmolarity did not adapt, we predicted that the nuclear volume *v_n_* would decrease with GIP, denoted *P*, as (more details in section II.A of the SI):

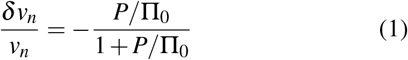

where Π_0_ is the intracellular nominal (P = 0 MPa) osmotic pressure and *P* corresponds to the surplus internal osmotic pressure above Π_0_. We fitted the nuclear volume data with 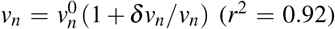 and obtained Π_0_ ~ 0.95 ± 0.05 MPa (dashed line Fig. 1c). We measured the osmotic pressure of the culture medium at 30°C to be Π_e_ ~ 0.63 MPa, leading to a nominal osmotic pressure difference between the cell interior and the cell exterior ΔΠ ~ 0.3 MPa, in agreement with values from the literature[24]. Since macromolecule concentration was increasing while cell volume remained constant, we predicted that macromolecular crowding would increase under GIP.

Changes in macromolecular crowding can be inferred by particle tracking microrheology[6]. We recently developed genetically-encoded multimeric (GEM) nanoparticles as highly efficient tracer particles for microrheology[6]. Introduction of a gene that encodes a self-assembling scaffold protein tagged with a fluorescent protein generates cells that constitutively contain tracer particles of defined sizes. In this study, we used GEMs of 20 nm (20nm-GEMs) and 40 nm (40-nm-GEMs) diameter. These particles probe the mesoscale, the length-scale of multimeric macromolecular assemblies such as RNA polymerase and ribosomes.

Using probes of various sizes, we found that the increase in cytoplasmic crowding under mechanical compression depended strongly on length-scale (Fig. 1d): the effective diffusion of larger particles such as mRNA (~ 80 nm diameter[25]), decreased far more than that of smaller particles such as 20nm-GEMs. We also found that the diffusion of a DNA locus decreased with GIP, probably as a consequence of decreased nuclear volume leading to increased nuclear crowding. Interestingly, the diffusivity of every tracer particle was compatible with an exponential decay similar to what was observed *in vitro* in [18], more apparent for larger mRNP particles (Fig. S3):

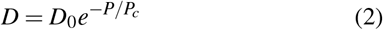

Where *D_0_* is the nominal diffusion of each particle, *P* is the growth induced pressure (GIP), and *P_c_* is the characteristic pressure of the exponential dependence on GIP for each particle. This exponential dependence of diffusion on GIP is theoretically predicted from the Doolittle relationship, as previously described[6] (see Modeling in SI). However, as above, this prediction only applies if osmolytes and macromolecules maintain a fixed, proportional concentration. We found that *P_c_* ∝ Π_*i*_/*ζ*, where *ζ* is a constant related to the interactions of the nanoparticle with its surroundings. Using osmotic perturbations to instantaneously modify crowding (Fig. S4), we were able to measure *P_c_* ~ 0.6 MPa for 40nm-GEMs. Using this value our theory predicts well our empirical data (solid black line in Fig. 1d is the prediction, red dots are the data).

Experimentally, we found that the characteristic pressure *P_c_* depended on particle size, and was inversely proportional to the probe size *s* (1/*P_c_* = *βs* where *β* is the proportionality constant, Fig. 1e). This inverse relation implies that the effective cytosolic diffusion for a particle of any size, *s* (in nm) is a power law function of the diffusion at 40 nm: *D_s_* ∝ *e*^−*βsP*^ = *e*^−*βsP*40/40*^ = (*e*^−*β*\*40\**P*^)^*s*/40^ ≈ D_40nm_^s/40^. Using this relationship, we can predict cytosolic diffusivity at any length-scale from the effective diffusion of 40nm-GEMs (*D*_40nm_).

Our data strongly suggest that confined growth leads to a concomitant increase in both internal osmotic pressure (leading to GIP and cell deformation) and macromolecular crowding (as evidenced by nuclear compaction and decreased nanoparticle diffusivity). Theory successfully predicts these observations if the increase in GIP and crowding are proportional. Another prediction of this proportional coupling is that relaxation of mechanical stress should lead to a cell volume increase proportional to GIP (see Modeling), *δv/v* = P/Π_0_; and for macromolecular crowding (and thus diffusivity) to reset to the nominal value without GIP. To test this prediction, we used a device in which GIP could be quickly relaxed Fig. S5). Consistent with our model, we observed a fast, fully reversible and predictable increase in cell volume, and recovery of GEM diffusion upon instantaneous relaxation of GIP (Fig. 1f). Together, this set of observations show that confined growth is compatible with a proportional increase in osmolyte and macromolecule concentration.

We next sought to investigate how GIP affects cell growth and protein production (which is dependent on the rates of multiple biochemical reactions). We first measured changes in cell number and chamber volume to estimate the cellular growth rate (Methods, Fig. S6). We observed that growth rate decreased roughly exponentially with GIP (Fig. 2a). To get insight into protein production, we used a fluorescent reporter assay. Protein production can take hours, raising the problem that GIP would continue to increase during the experiment if growth continued. To avoid this issue, we expressed the mCherry fluorescence protein from the *ADH2* promoter (P_ADH2_-mCherry) as our model system. The *ADH2* promoter is activated by glucose starvation, a condition that also arrests cell growth[26]. Thus, we could grow cells to develop a defined amount of GIP and then induce *P_ADH2_-mCherry* by withdrawal of glucose (osmotically balancing with sorbitol) at a range of GIP values. In this way, we could infer how protein expression, at least of this model gene, was affected by GIP. We observed that the induction of the fluorescence signal was slower under GIP than in the control (Fig. 2b).

**Figure 2.**
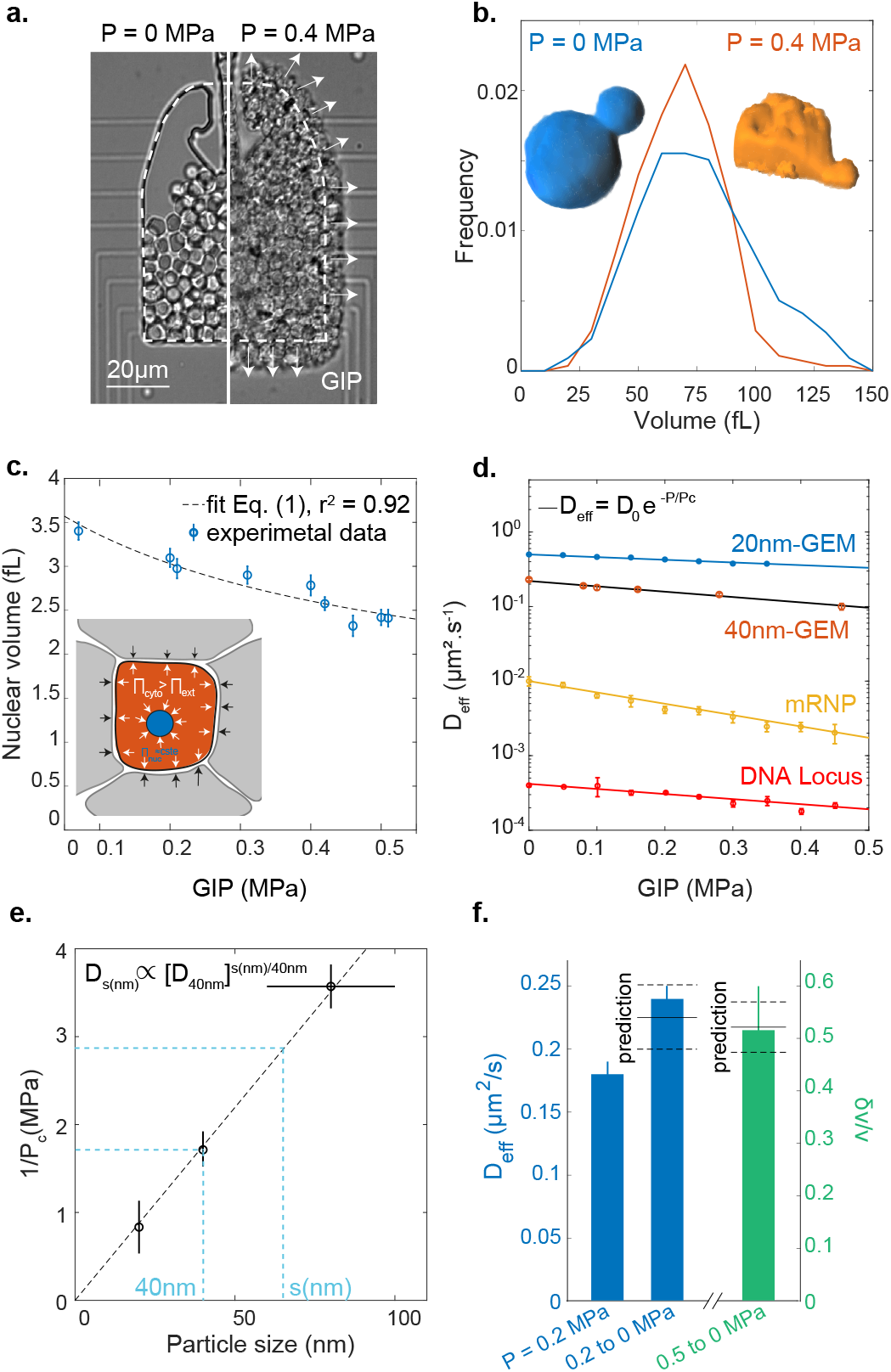
Confinement decreases growth and protein production rates. **a.** Growth rate decays roughly exponentially with GIP. **b.** Representative images from protein production reporter system. A reporter gene consisting of the mCherry fluorescent protein under the control of the *ADH2* promoter was integrated at the endogenous locus; glucose starvation induces the gene. After 7 h of induction, we observe stronger induction for control (no pressure, left) condition than under GIP (right). **c.** Single cell fluorescence intensities were fitted with a quadratic function (see Method) to extract an effective expression rate *k*_exp_ at various values of GIP. Representative curves for a single cell is shown with fitting; multiple single cell traces are shown inset. **d.** Protein expression rate decreases roughly exponentially with GIP. In all data, values are mean ± standard error of the mean, over n ≥ 100 cells in N ≥ 3 independent biological replicates.

This experimental strategy enabled us to extract single-cell *P_ADH2_-mCherry* fluorescence intensity curves. We observed that, after an initial time delay, which could be associated with sensing of carbon starvation or promoter remodeling, fluorescence intensity increased with time, and that this rate of increase was lower in compressed cells. We developed a mathematical model of transcription followed by translation to quantify induction of fluorescence. Our model predicted that protein concentration should increase quadratically with time at short time scale, with an effective rate *k*_exp_ that is the product of the transcription rate (*k*_m_) and the translation rate (*k*_p_): *k_exp_* = *k_m_* * *k_p_* (see Modeling). Although *k_exp_* was not *stricto sensu* a rate, but rather the product of two rates, we refer to it as a single effective rate hereafter, for the sake of simplicity.

Our simple model yielded an excellent fit to the experimental data (Fig. 2c), and enabled us to extract both the time delay and *k_exp_*. We observed that the time delay was progressively shorter with GIP (Fig. S7). It has previously been shown that cells in the G1 phase of the cell cycle respond more rapidly to stress[26], and our previous studies showed that *S. cerevisiae* arrests in G1 in response to GIP[5, 27]. Therefore, accumulation of cells in G1 could explain this reduced lag-time under GIP. We found that *k_exp_* decreased roughly exponentially with GIP (Fig. 2d), with a similar dependence as the growth rate decay (about 60% decrease at *P* = 0.3 MPa in both cases). We confirmed the reduction of protein production rate on the expression of a constitutively active *P*_HIS3_-*GFP* construct, suggesting that our results were not specific to the use of *P*_ADH2_-*mCherry* (Fig. S8).

Our microrheology data and nuclear compression demonstrated that macromolecular crowding increased under GIP. We hypothesized that this crowding could limit protein expression rate and ultimately growth itself. This feedback could be physical as a result of decreases in the rate of diffusion-limited processes, with no need for specific signaling pathways. To test this idea, we set out to perturb molecular crowding by orthogonal means, using osmotic compression.

Increasing external osmotic pressure, for example by addition of sorbitol to the media, leads to water efflux from cells, reducing cell volume and increasing the concentration of biomolecules within the cell. This osmotic compression has previously been shown to increase macromolecular crowding[17]. Wild-type cells rapidly adapt to these perturbations through the osmotic response pathway, controlled by the *Hog1p* kinase, which increases production of intracellular glycerol to counteract the increased external osmotic pressure. The rapid recovery from osmotic perturbation makes it difficult to interpret long-term experiments in wild type cells. Furthermore, intracellular viscosity is affected by glycerol accumulation, making rheological measurements hard to interpret. To avoid these issues, we used *hog1*Δ mutant cells, which cannot rapidly respond to acute osmotic stress[28]. We and others find that *hog1*Δ cells can still expand and grow, even in the presence of increased concentrations of external osmolytes[29] indicating that there is still baseline generation of internal osmolytes[30]. Therefore, at least two mechanisms generate intracellular osmolytes. First, a basal mechanism constitutively generates the osmotic pressure that cells require to expand and grow; we predict that this basal osmolyte production is coupled to the rate of macromolecule biosynthesis, but is not increased to allow cells to adapt to osmotic shock. Second, an acute stress response, dependent on *Hog1p*, allows cells to adapt to changes in external osmotic pressure.

We performed laser ablation experiments to confirm that *hog1*Δ cells still maintain internal osmotic pressure, even after osmotic compression with 1M sorbitol (Fig. S9). This experiment confirms that osmotic compression is a useful orthogonal approach that allows us to increase crowding at all length-scales, very similar to GIP. By using *hog1*Δ cells, we are able to maintain increased crowding for sufficient time to assess growth and protein expression rates.

GIP and osmotic compression are orthogonal means of increasing molecular crowding and activate distinct stress response pathways. Furthermore, the osmotic stress response is largely abrogated in *hog1*Δ cells[28]. We found that *k*_exp_ of *P*_ADH2_-*mCherry* decreased with osmotic compression (Fig. 3a). If the effective expression rate (*k*_exp_) of *P*_ADH2_-*mCherry* is modulated by macromolecular crowding, then *k*_exp_ should display the same relationship to the effective diffusion of 40nm-GEMs (*D*_40nm_) under both GIP and osmotic compression. Indeed, we observed the same dependence in both conditions (Fig. 3b) supporting the hypothesis that macromolecular crowding limits protein expression.

**Figure 3.**
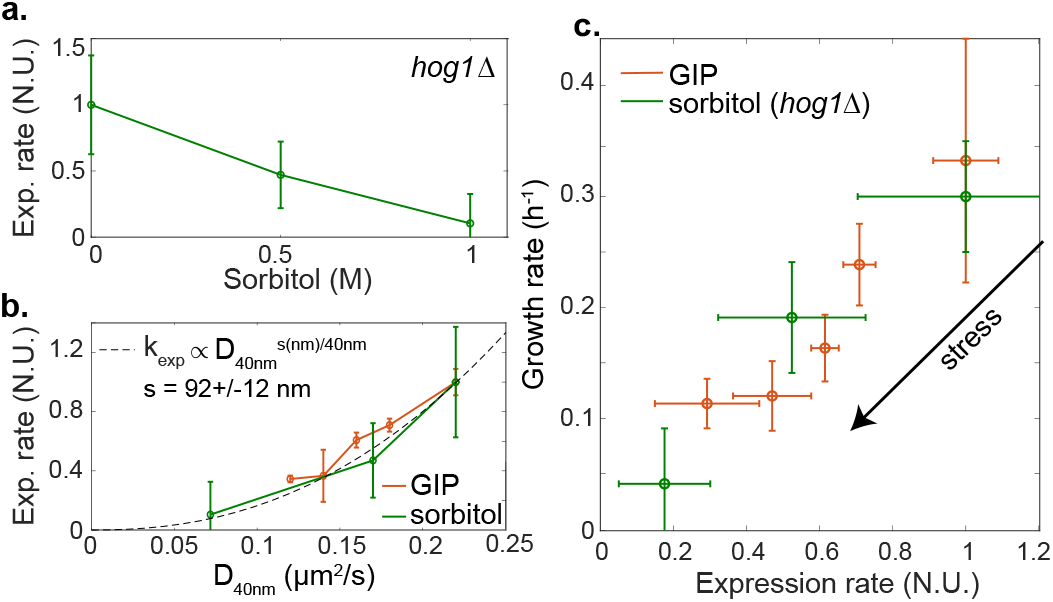
Protein production and growth are diffusion limited processes. **a.** Effective expression rate (*k*_exp_) of *P*_ADH2_-*mCherry* decreases under osmotic stress in *hog1*Δ cells. **b.** *k*_exp_ was fitted by a power-law function of the effective diffusion of 40nm-GEMs. **c.**Growth rate is proportional to protein production rate for *hog1*Δ cells under osmotic stress and *WT* cells under GIP. In all data, values are mean ± standard error of the mean, over N ≥ 3 independent biological replicates.

Our results are consistent with effective protein expression rate being diffusion-limited at a certain unknown length-scale, *s*. We found that the relationship between effective diffusion and particles diameter was a power law (Fig 1e). If crowding decreases P_ADH2_-*mCherry* production by inhibiting diffusion of a rate-limiting particle, effective expression rate should be a power law function of *D*_40nm_ with an exponent that is the ratio of the particle size, *s* (in nm), divided by the size of 40nm-GEMs (i.e. 40 nm):

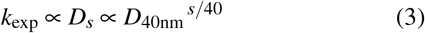

Indeed, we observed this power-law dependence, with an exponent that suggests that expression was limited by the diffusion of particles of a characteristic size s ~ 90 nm (Fig 3b). This mesoscale length-scale corresponds to many biological entities, for example trafficking vesicles and mRNA ribonucleoprotein particles[25, 31, 32] (both ~ 100 nm).

We next investigated the hypothesis that growth rate is mainly limited by protein expression. We plotted growth rate as a function of the effective expression rate of *P_ADH2_-mCherry* and found that the two rates were roughly proportional. Note, this model gene is not limiting growth-rate, *ADH2* is not expressed in the presence of glucose. Nevertheless, the fundamental processes required for its expression (e.g. transcription by RNA polymerase II and translation by ribosomes) are shared by all proteins. Interestingly, we observed that the same relationship held for both osmotic compression of *hog1*Δ cells and *WT* cells under GIP (Fig. 3c). Even osmotically compressed, *hog1*Δ cells are still able to grow. The fact that growth rate similarly decreases with protein production under both osmotic stress and GIP indicates that similar limiting mechanisms could be at play.

Taking all of our results together, we developed a model of confined growth, with all parameters experimentally determined, allowing us to predict protein production and cell growth in confined conditions. The model derivation and parameterization are detailed in the Supplementary Information.

Our data support a central hypothesis that osmolyte and macromolecule production rates are tightly coupled. Exactly how this balance of rates is achieved remains unknown and is a longstanding fundamental question, but as a consequence, in the absence of confinement, cells grow and accumulate biomass while maintaining a constant level of macromolecular crowding. The accumulation of osmolytes increases osmotic pressure. The mechanical balance between osmotic pressure and the elastic properties of the cell wall in turn defines the turgor pressure[33]. This turgor pressure enables the cell wall to expand through a process of hydrolysis and insertion of new cell wall material[30, 34]. We posit that insertion of cell wall material is only possible when the turgor pressure resulting from osmolyte accumulation is above a fixed value.

If the effective elasticity of the cell wall, encompassing the various mechanical parameters such as its Young modulus and Poisson ratio or thickness, were to increase (i.e. require more force to be deformed), a higher pressure-difference, and thus more osmolytes, would be required to achieve expansion. This is also the case during confined growth where the surroundings mechanically resist cell growth. Confined growth leads to an effective increase in the elasticity around the cell which then physically limits cell wall expansion (Fig. 4a). In our experiments, when cells fill the confining chamber and start to distort one another and the chamber walls, they experience an effective surrounding elasticity, *E*_eq_. When cells grow by *δv*,they need to accumulate more osmolytes to expand the cell wall, resulting in an increased internal pressure, which is the product of the surrounding effective elasticity and the volume change: *E*_eq_ * *δv/v*. This value is the growth-induced pressure (GIP). Based on our central hypothesis that the accumulation of osmolytes is proportionally coupled to the accumulation of macromolecular biomass, the decreased expansion rate will lead to increased crowding. This increase in crowding then feeds back onto both protein and osmolyte production, which further reduces the cell expansion rate (Fig 4a).

**Figure 4.**
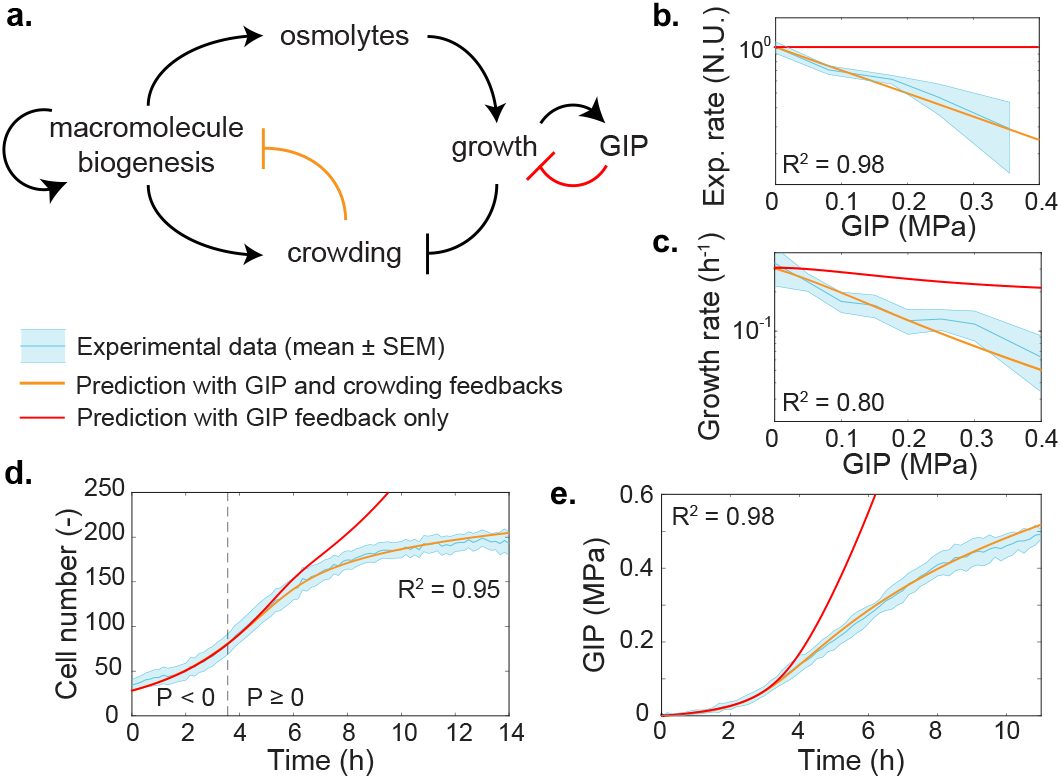
A physical feedback model, in which crowding limits protein production, predicts the dynamics of confined cell growth. **a.** Schematic of the model. After confluence is reached, growth induced pressure (GIP) increases as a function of cell volume change and the effective elasticity of the surrounding cells and PDMS chamber walls. Cells must accumulate more osmolytes to grow in the face of increasing effective elasticity, therefore volume expansion is inhibited. Macromolecule biogenesis is proportionally coupled to osmolyte production and so intracellular crowding increases. Increased mesoscale crowding feeds back onto many processes including the processes associated with macromolecule biogenesis itself, thereby limiting growth. **b-e.** Predictions of the dependence of various observables on GIP from the model. All parameters are experimentally determined. Predictions are shown for: effective protein expression rate (*k*_exp_, **b**); growth rate (**c**); cell number (**d**); and GIP (**e**). In all plots, the thick orange line represents the model prediction. The thick red line represents the prediction of the model without any physical (crowding) feedback on biomass production. The dashed line represents the onset of confluence and GIP buildup. The *R^2^* value indicates the square difference of the model against the data. Values are mean ± standard error of the mean; N ≥ 3 independent biological replicates.

We calibrated the parameters related to our confining growth model, including the value of the turgor pressure, using laser ablation, transmission electron microscopy and atomic force microscopy (Supplementary Information).

Our experimentally calibrated model accurately predicted the dependence of protein production and cell growth rate on pressure, as well as the dynamics of confined cell proliferation and GIP buildup, without any fitting of free parameters (thick orange lines in Figs. 4b-d). This remarkable predictive power supports our simple model: Growth is initially limited by the surrounding elastic environment, which forces the cell to increase internal osmolarity. Osmolyte production is directly coupled to macromolecular biosynthesis thus leading to mesoscale crowding. High mesoscale intracellular crowding then physically inhibits reactions through diffusion-limited processes. Our model shows that most of the observed decrease in growth rate can be explained by this physical feedback, without the need to evoke any other mechanism.

We also investigated the predictions of our model if we removed the physical feedback (thick red lines in Figs. 4b-d). In this case, GIP and cell number would rise much more quickly than experimentally observed. Growth would still ultimately decrease, due to the increasing mechanical barrier to cell expansion, but much more slowly than observed because the rate of osmolyte production would not be limited. In this case, crowding would also rise quickly, and crowders in the cell would approach the maximum random close packing fraction much sooner. We speculate that the physical feedback of crowding on biosynthesis is adaptive, as it delays and attenuates macromolecular overcrowding, which could allow more time for stress responses to more efficiently activate. Which step of protein biosynthesis is limited by crowding is however unknown and requires a separate investigation.

An intriguing question is why cells have not evolved adaptive mechanisms to change the relative rates of macromolecular biosynthesis and osmolyte production to prevent overcrowding of the cell. The osmotic stress response is an adaptive mechanism of this type. However, we observed that GIP in *hog1*Δ mutants, which are defective for the osmotic stress pathway, was similar to that in wild-type cells (Fig. S10). A key difference between GIP and osmotic shock is the effect on turgor. The activation of the osmoadaptive *HOG1* pathway in *S. cerevisiae* is linked with a loss of turgor[35]. However, our results suggest that turgor does not decrease during GIP, and in fact increases due to the effective elasticity of the surroundings effectively acting like a thicker cell wall. Increased turgor actually triggers the hypo-osmotic stress response, which decreases intracellular osmolarity and subsequently cell volume[36]. However, this would be counterproductive during confined growth as reduced cell volume would further increase crowding. Indeed, pathways related to the response to both hyper- and hypo-osmotic stress are triggered by GIP. These pathways, which together constitute the SCWISh network[16], are important for cell survival under GIP, but they do not appear to change the coupling between osmolyte and macromolecule biosynthesis. Perhaps the feedback between mesoscale crowding and growth is useful: diffusion is affected with a strong size-dependence, mainly limiting reactions at the mesoscale (≥ 10 nm diameter). It is intriguing that many stress response proteins are relatively small. Therefore, upon developing strong growth-induced pressure, growth will stall, but stress-response pathways can continue to operate.

Stress-response signaling pathways vary extensively between organisms. In contrast, high macromolecular crowding is a fundamental property of all life forms[37]. Our results suggest that a primordial biophysical feedback mechanism arises directly from the physical properties of cells. This feedback could be essential for multicellular proliferation, and its deregulation important in the context of some pathologies. Solid cancer cells in particular, in contrast with normal cells, acquire the capacity to proliferate under confinement and build up GIP, suggesting that genetic alterations, or chemical environmental modifications, can impact the ability to proliferate under confinement.

## Acknowledgments

We thank E. Kassianidou for initial help with laser ablation experiments. We thank NYULH DART Microscopy Laboratory Alice Liang, Chris Petzold and Kristen Dancel-Manning for consultation and assistance with TEM work; this core is partially funded by NYU Cancer Center Support Grant NIH/NCI P30CA016087. The technological realizations and associated research works were partly supported by the French RENATECH network (MD). LJH was funded by NIH grants R01 GM132447 and R37 CA240765. We thank E. Rojas for fruitful discussions. LJH and MD thank the FACE foundation for travel support.

## Author Contributions

BA and MD designed and performed the experiments and data analysis; CCD and ED performed the AFM experiments; LJH designed the strains used in the study; BA, LJH and MD wrote the manuscript.

## Data availability

Source data are available for this paper. All other data that support the plots within this paper and other findings of this study are available from the corresponding author upon reasonable request.

